# Predatory behaviour as a personality trait in a wild fish population

**DOI:** 10.1101/2020.06.19.161968

**Authors:** Andrew W. Szopa-Comley, Callum Duffield, Indar W. Ramnarine, Christos C. Ioannou

## Abstract

Consistent inter-individual differences in behaviour (i.e. animal personality variation) can influence a range of ecological and evolutionary processes, including predation. Variation between individual predators in commonly measured personality traits, such as boldness and activity, has previously been linked to encounter rates with their prey. Given the strong selection on predators to respond to prey, individual predators may also vary consistently in their response to prey in a manner that is specific to the context of predation. By studying wild piscivorous fish (pike cichlids, *Crenicichla frenata*) in their natural environment using experimental presentations of prey and control stimuli, we show that individual predators differ consistently in the amount of time spent near prey. Crucially, these differences were not explained by the behaviour of the same individuals in control presentations (the same apparatus lacking prey), suggesting that variation in the response to prey reflects a ‘predator personality trait’ which is independent from other individual traits (body size, boldness and/or neophobia) and environmental factors. Pike cichlids which spent more time near prey also attacked prey at a higher rate. These findings imply that the risk posed by individual predators cannot always be adequately predicted from typically studied axes of personality variation.

## Introduction

Through their direct effect on prey abundance (Paine 1966) and non-lethal impact on prey physiology and behaviour (Beckerman et al. 1997; Lima 1998), predators exert a strong influence on the structure and composition of ecological communities (Schmitz et al. 2004). An extensive body of research has explored how prey adjust their behaviour in response to changes in predation risk (Lima and Dill 1990), and has revealed how the mere presence of nearby predators can shape prey population dynamics and the abundance of resources at lower trophic levels (Preisser et al. 2005; Peckarsky et al. 2008; Suraci et al. 2016). Predators, by contrast, are often viewed as being behaviourally unresponsive, posing a fixed and uniform level of risk to prey (Lima 2002). Increasingly this simplifying assumption is at odds with the evidence for widespread consistent inter-individual differences in behaviour (also known as personality variation) within natural populations (Bell et al. 2009). As the effects of variation within species on ecological processes can often equal or outweigh the impact of differences between species (Des Roches et al. 2018), determining how individual predators differ in their behaviour is key to understanding their wider effects (Ioannou et al. 2008; Okuyama, 2008; Start and Gilbert 2017; Michalko and Řežucha, 2018; Rhoades et al. 2019).

Empirical research has shown that inter-individual behavioural differences can influence numerous ecological and evolutionary processes (Dall et al. 2012; Sih et al. 2012), ranging from dispersal (Cote et al. 2010) to pair bonding (Firth et al. 2018). However, most studies have focused on a limited number of traits, particularly boldness, exploration, activity, aggressiveness and sociability (Réale et al. 2007), which are not necessarily the most ecologically relevant axes of variation (Koski 2014). Although a wide variety of other behaviours are known to be individually repeatable (reviewed in Bell et al. 2009), there have been few tests examining whether inter-individual variation in these behavioural traits is separate from, or correlated with, frequently measured axes of variation. If commonly studied personality traits are not strongly correlated with other repeatable behaviours that have greater ecological or evolutionary relevance, this has implications for both how personality traits affect ecological and evolutionary processes, as well as the selection imposed on different personality traits. Answering this question may be particularly important for widespread ecological processes like predation, which is almost ubiquitous in animals across a diverse range of taxa and habitats.

The majority of studies exploring the consequences of personality differences for predator-prey interactions have concentrated on variation in activity levels or boldness (Bell and Sih 2007), behaviours which reflect the degree to which individuals prioritise gaining resources over risk avoidance (Smith and Blumstein 2008). Bolder or more active prey are often more susceptible to predation (Ballew et al. 2017; Hulthén et al. 2017), although this does not necessarily result in positive correlations between boldness and survival (Moiron et al. 2020). The link between these traits and individual movement patterns also suggests that bolder or more active predators are also more likely to encounter prey (Spiegel et al. 2015). Consistent with the expected effect of boldness on encounter rates, the interacting effects of prey and predator behavioural types have been shown to determine prey survival (Pruitt et al. 2012; Chang et al., 2017). In seabird populations, boldness also predicts inter-individual variation in how individual predators search for prey (Patrick et al. 2017). As well as affecting encounter rates with prey, personality variation influences the rate of prey consumption (Toscano and Griffen 2014), and individual predators have also been shown to differ consistently in the time taken to detect, respond to or capture prey (McGhee et al. 2013; Szopa-Comley et al., 2020; MacGregor et al., 2020).

Quantifying behavioural variation in the wild avoids artefacts which can arise when behaviour is expressed in laboratory conditions (Niemelä and Dingemanse 2014). Predator personality variation will also be most relevant to the risk prey experience when prey are repeatedly exposed to the same individual predators. This context arises in populations of the Trinidadian guppy (*Poecilia reticulata*) in their natural habitats, where guppies can be confined to the same natural river pools during the dry season as pike cichlids (*Crenicichla frenata*), their main predator (Magurran 2005; Botham et al. 2006). Rather than engaging in lengthy pursuits, pike cichlids typically track their prey visually before attacking in a rapid burst once they have approached within close proximity (Walker et al., 2005; Heathcote et al., 2020). In this study, we investigated inter-individual variation in predator behaviour by repeatedly presenting stimulus shoals of guppies *in situ* (the prey treatment) over multiple days across a series of 16 discrete natural river pools, and recorded the time spent near the stimulus by individually-identified pike cichlids (Fig. 1). Similar methods of quantifying predation risk have also been shown to correlate with anti-predator behaviour in this system (Croft et al. 2006). Importantly, we repeatedly presented the same apparatus without prey as a control treatment in each pool. The response of individual predators to the control should reflect variation in the individual traits or environmental factors which are not specific to prey, including personality variation in boldness and neophobia (during the initial presentation, the (empty) stimulus was entirely novel). For example, whether or not individual pike cichlids were recorded approaching the control stimulus should indicate their response to novel features within their environment. By comparing the predator’s behaviour in the prey and control treatments, we were therefore able to isolate inter-individual variation in predatory behaviour (i.e. the response to prey) from factors that are not specific to prey.

**Figure 1:**
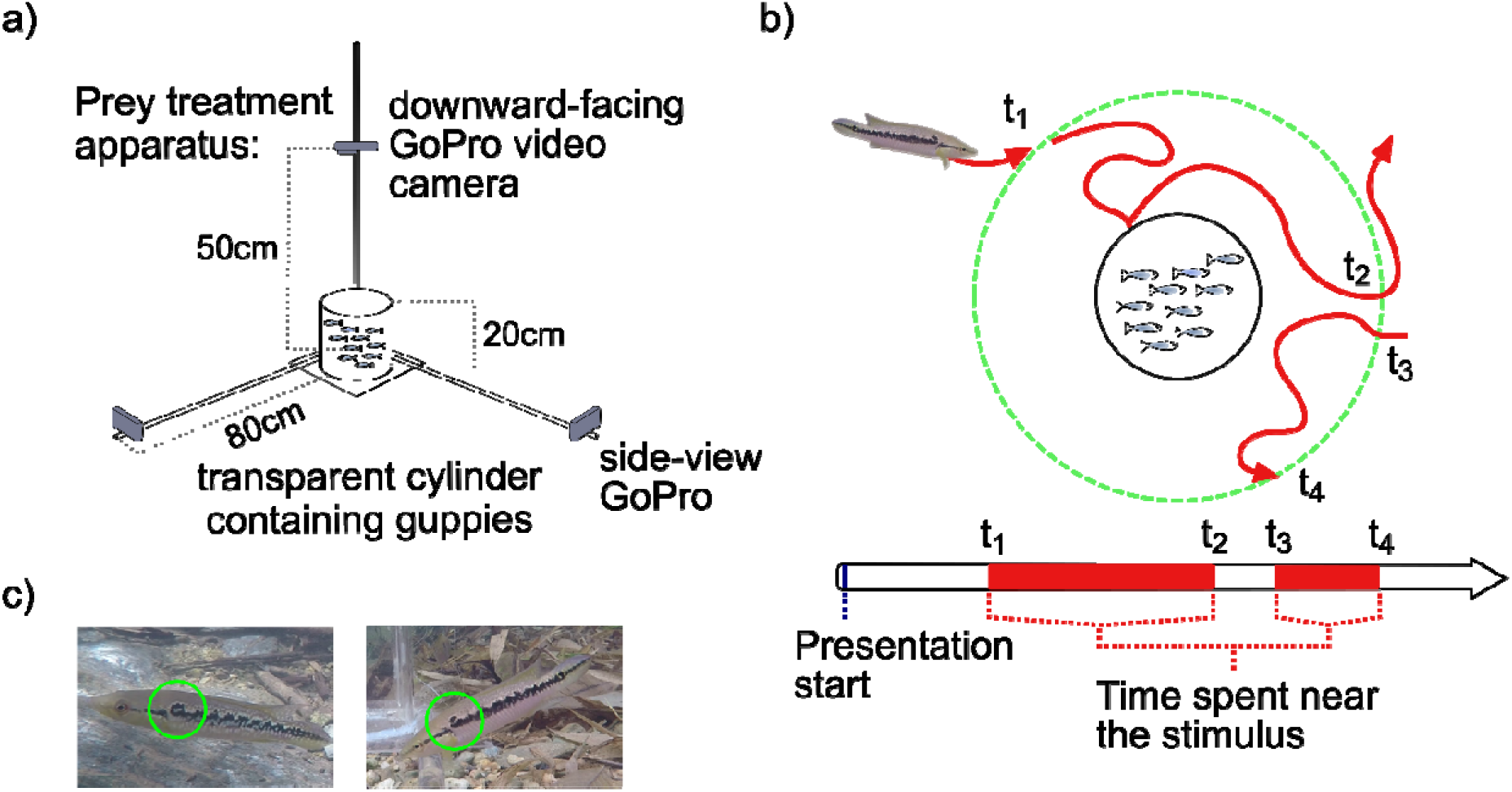
Quantifying inter-pool and inter-individual variation in predatory behaviour. a) For each river pool, the apparatus used in the study was deployed in one of two treatments: in the prey treatment, 10 female guppies were placed within the cylinder (diagram not to scale), or in an otherwise identical control apparatus without guppies (not shown). b) Depending on the level of the analysis (inter-pool or inter-individual differences), the time spent near the stimulus was defined as the total amount of time in which any pike cichlid (or a specific individual) was present within a zone surrounding the stimulus during the 30-minute presentation period. The green dashed line (approximately 24cm from the outer edge of the cylinder) represents the outer limit of the zone surrounding the stimulus, viewed from above. The total time spent near the stimulus in each presentation is the sum of the time difference between t_1_ and t_2_ plus the time spent within the zone during subsequent approaches (denoted by t_3_ and t_4_, etc.). c) Individual pike cichlids were identified through differences in humeral spot patterns (highlighted by green circles), using still images recorded by the side-view GoPro video cameras.

## Methods

### Study site

We used a series of discrete natural river pools in the Lopinot valley, Trinidad (Appendix: Table A1). Pools in this river are characterised by deeper areas of water (>0.5m in depth), bounded by shallower sections containing riffles, rocks, boulders and small waterfalls which restrict the movements of fish. Other, although minor, predators known to be present in this river include blue acara cichlids (*Aequidens pulcher*), two-spot astyanax (*Astyanax bimaculatus*) and wolf-fish (*Hoplias malabaricus*) (Magurran 2005).

### Apparatus

The apparatus consisted of a transparent acrylic plastic cylinder (diameter: 15cm, height: 20cm, wall thickness: 3mm) attached to a square base and covered by a perforated lid. Approaches to the apparatus were recorded using Go Pro video cameras attached to the ends of three clear acrylic plastic rods (diameter: 3cm, length: 80cm) secured to the base of the apparatus, with one rod extending vertically and two identical rods projecting horizontally at right angles to one another (Fig. 1a). This arrangement allowed footage of approaching fish to be captured from above (Hero 3+ video camera, frame rate: 25 frames per second, resolution: 960p) and from the side (two GoPro Hero 5 video cameras, frame rate: 30 frames per second, resolution: 2.7k; diagonal field of view for both camera models: 133.6°). The apparatus was manoeuvred into position from the edge of each pool with minimal disturbance using transparent monofilament attached to the base.

### Experimental procedure

The study was designed to quantify inter-individual differences in the behavioural response of predators to their prey by repeatedly presenting a stimulus prey shoal (the prey treatment, consisting of 10 female guppies placed within the cylinder of the apparatus) to free-swimming predatory fish in their natural environment. Control presentations of an otherwise identical apparatus without guppies were also conducted in the same locations to measure variation in the individual traits of the same predator individuals (such as personality variation in boldness and neophobia) which have been shown to influence encounter rates with prey. With this approach, it was then possible to test for the existence of a separate predator personality trait by examining individual responses to prey and responses of the same individuals when prey were absent.

Experiments were carried out over a six-week period from March to May 2017. At the start of each 30-minute presentation period, the apparatus was placed in the same location and orientation within each pool. For each river pool, the apparatus used in the study was deployed in one of the two treatments in six separate presentations (once per day): three prey treatment presentations and three control presentations. Presentations took place between 0730 and 1130 over a period of six consecutive days. Within each six-day period, control presentations of the entirely novel apparatus always took place on the first day. Guppies used in prey treatment presentations were caught from a single pool in the same river using a seine net (the stimulus was not deployed in this pool). In each prey treatment presentation, 10 female guppies of a similar size were selected haphazardly from a number collected at the start of each day. The order in which the two treatments were assigned to the remaining five presentations per pool was randomised, but for logistical reasons, the three pools tested over the same six-day period shared the same presentation order. As incident light levels may affect pike cichlids’ ability to detect prey, pool canopy openness was also measured after completion of the experiment by averaging measurements taken with a spherical densiometer in all four cardinal directions at three points along at the pool’s upstream-downstream axis: the upstream end, midpoint and downstream end.

### Video analysis

Data on the behavioural response of pike cichlids in each pool were extracted at two levels: pool-level data on the time spent near the stimulus by any predator individual over the course of the 30-minute presentation period, and individual-level data on the time spent near the stimulus by individual pike cichlids. In both the pool- and individual-level analyses, the time spent near the stimulus was defined as the total amount of time in which any pike cichlid (or a specific individual) was present within a zone surrounding the stimulus during the 30-minute presentation period (the zone extended to approximately 24cm from the outer edge of the cylinder; Fig. 1b). The number of pike cichlids in each pool was estimated using still images recorded using the side-view cameras (fish which could not be conclusively identified were not included in this total). The standard body length of individual pike cichlids (median approximate standard body length: 9.8 cm, inter-quartile range: 2.9 cm) was quantified in ImageJ (version 1.46r) using still images obtained from the downward-facing video camera (Fig. 1c) and comparing fish-length measurements in pixels to an object of known length (the stimulus cylinder viewed from above).

### Statistical analysis

All analyses were carried out using R v. 3.3.2 (R Development Core Team 2019), and all LMMs (linear mixed effects models) and GLMMs (generalised linear mixed effects models) were fitted with the lme4 package. Results and further details for all models (including response variables, explanatory variables included as fixed effects, random effects and sample sizes for each model) are given in Appendix: Table A2. Prior to the analysis, all continous explanatory variables were standardised by subtracting the mean and dividing by the standard deviation. In each model, *P*-values for fixed effects were derived from likelihood ratio tests comparing the full model with a reduced model lacking the variable in question. Model assumptions were verified by plotting residuals versus fitted values, using the DHARMa R package for GLMMs (Hartig 2019).

To quantify inter-individual differences in predatory behaviour, estimates of adjusted repeatabilities and inter-individual variances were obtained using the rptR package (Stoffel et al. 2017), which utilises a mixed model framework (Nakagawa and Schielzeth 2010), allowing experimental (presentation number and time of day) and environmental variables (canopy openness and the estimated number of pike cichlids in each pool) to be included as fixed effects. Adjusted repeatabilities can thus be interpreted as a standardised measure of inter-pool or inter-individual variation after potentially confounding variables have been controlled for. LMMs were used to analyse pool-level data and Poisson GLMMs were used to analyse individual-level data, which also included an approximate measure of pike cichlids’ standard body length as an additional fixed effect to control for predator body size. Pool identity was included as a random intercept in models used to analyse pool-level data. Models used to analyse individual-level data included both pool and individual identity as nested random intercepts. Statistical significance of repeatability estimates was assessed using both *P*-values (obtained through likelihood ratio tests) and overlap of the 95% confidence intervals with zero (computed via parametric bootstrapping). Data on the time spent near the stimulus was censored at a maximum of 30 minutes corresponding to the duration of a presentation, but no fish spent the maximum amount of time near the stimulus. Unless otherwise stated, instances when fish spent zero time near the stimulus were disregarded in the analysis in order to avoid influencing estimates of within-pool or within-individual variation and thus affecting repeatabilities (Stamps et al. 2012).

Social interactions between predators within the same pool could potentially generate feedbacks which magnify (via differentiation) or suppress (via conformity) inter-individual differences in behaviour. To test for these possibilities, we conducted two types of randomisation simulations based on data for predators which approached the prey treatment stimulus in multiple presentations (44 individuals), following the methodology outlined in Ioannou et al. (2017). In the first randomisation, the mean observed pool-level diversity in the response to prey between predators across all pools was compared to the distribution of this statistic produced when the set of observations corresponding to an individual predator was randomly exchanged between pools. The aim of this approach was to determine whether the response of individual predators to prey was dependent on the other individuals present within the same pool, or whether being in a particular pool is statistically unimportant. Pool-level diversity in the response to prey was quantified as the coefficient of variation (COV) in the time spent near the prey treatment stimulus. The second randomisation examined the relationship between the time spent near the stimulus by two individuals (‘predator 1’ and ‘predator 2’) which were randomly selected (without replacement) from the same pool. To enable direct comparison of the response to prey by ‘predator 1’ and ‘predator 2’, observations were selected from the same presentation per pair. Across multiple pools, a negative slope would be expected from behavioural differentiation between the two individuals (if one predator spends a long time near the stimulus, the other does not), whereas a positive slope would be consistent with conformity (both ‘predator 1’ and ‘predator 2’ spend a similar amount of time near the stimulus). The slope of the relationship between time spent near the stimulus by ‘predator 1’ and ‘predator 2’ was estimated using a quasi-Poisson generalised linear model, controlling for variation between pairs in presentation number, time of day, canopy cover and the estimated number of pike cichlids in each pool by including these variables as main effects. Both randomisation procedures were run with 10,000 iterations.

### Ethical note

Ethical approval for all procedures was obtained through the University of Bristol (project number UIN UB/17/006). The approach used in this study allowed the responses of free-swimming pike cichlid predators to prey to be measured, whilst also protecting prey from harm, through the presence of a physical barrier separating predators from prey. Guppies were collected from a population in the Lopinot valley which naturally coexists with pike cichlids, and after being used in the study, were returned to a pool in the same river.

## Results

### Responses to the experimental apparatus

A total of 69 individual pike cichlids were observed approaching the stimulus during at least one 30-minute presentation period (Appendix: Table A1). Compared to their behaviour in control presentations, pike cichlids spent more time near the stimulus when prey were present (Poisson generalised linear mixed effects model (GLMM), *N*_obs_ (no. of observations) = 211, *N*_ind_ (no. of individuals) = 69: χ^2^_1_ = 63.2, *P* < 0.001; Appendix: Fig. A1 and Table A2, model 1). In addition, attacks on the stimulus, defined as a fast, directed movement towards the apparatus (Appendix: Video A1) were only observed in the prey treatment. Pike cichlids which spent more time near the stimulus were more likely to attack the stimulus during the same presentation (binomial GLMM, *N*_obs_ = 133, *N*_ind_ = 68: χ^2^_1_ = 41.1, *P* < 0.001; Fig. 2; Appendix: Table A2, model 3), and made more attacks during the first 30 seconds that they spent near the stimulus (Poisson GLMM, *N*_obs_ = 123, *N*_ind_ = 63: χ^2^_1_ = 6.83, *P* = 0.009; Fig. 2; Appendix: Table A2, model 5).

**Figure 2:**
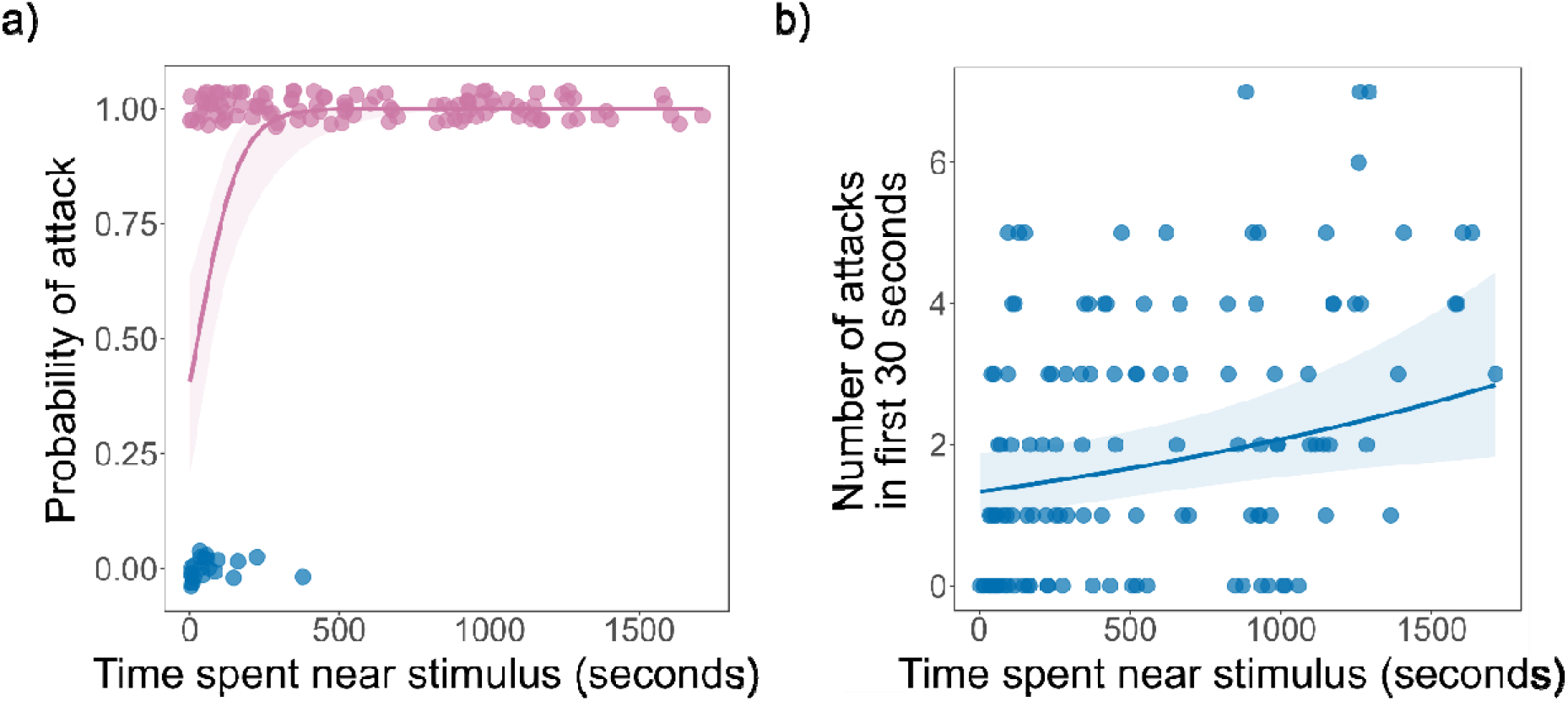
Relationship between the time spent near the stimulus and attack behaviour. a) Probability of attack as a function of the time spent near the stimulus. Jitter has been added to the raw data to allow overlapping points to be visualised. b) Relationship between the number of attacks made by pike cichlids during the first 30 seconds they spent near the stimulus and the total time spent near the stimulus during each presentation (this analysis was limited to pike cichlids which spent a minimum of 30 seconds near the stimulus). In both a) and b), curves represent the predicted response from a generalised linear mixed-effects model (GLMM): a binomial GLMM featuring attacks on the stimulus as a binary response variable in a) and a Poisson GLMM in b) (see Appendix: Table A2, models 3 and 5 respectively). Model predictions were obtained by holding all other fixed effects constant at their mean values. Shading indicates 95% confidence intervals surrounding the predicted response.

### Local variation in predation risk between river pools

From the prey’s perspective, the repeatability of predation risk in their local habitat (pools in our study) will be more relevant to the risk they experience than repeatable differences between individual predators. We therefore quantified inter-pool variation in the time spent near the stimulus by any pike cichlid in a given pool by estimating adjusted repeatabilities (Nakagawa and Schielzeth 2010), controlling for variation arising from the experimental design (time of day and presentation number) and environmental differences between the pools (the estimated number of pike cichlids in each pool and canopy openness). Significant repeatability at the level of the pool was evident during prey treatment presentations, even when controlling for the estimated number of pike cichlids, i.e. predator density, in each pool (*R*_pool_ = 0.476, 95% confidence intervals: 0.093 – 0.759, *P* = 0.009, *N*_obs_ = 45, *N*_pool_ (no. of pools) = 16; Fig. 3; Appendix: Table A3). The time spent near the stimulus by pike cichlids was not repeatable during control presentations without prey (*R*_pool_ = 0, 95% confidence intervals: 0 – 0.417, *P* = 1, *N*_obs_ = 35, *N*_pool_ = 15; Fig. 3; Appendix: Table A3).

**Figure 3:**
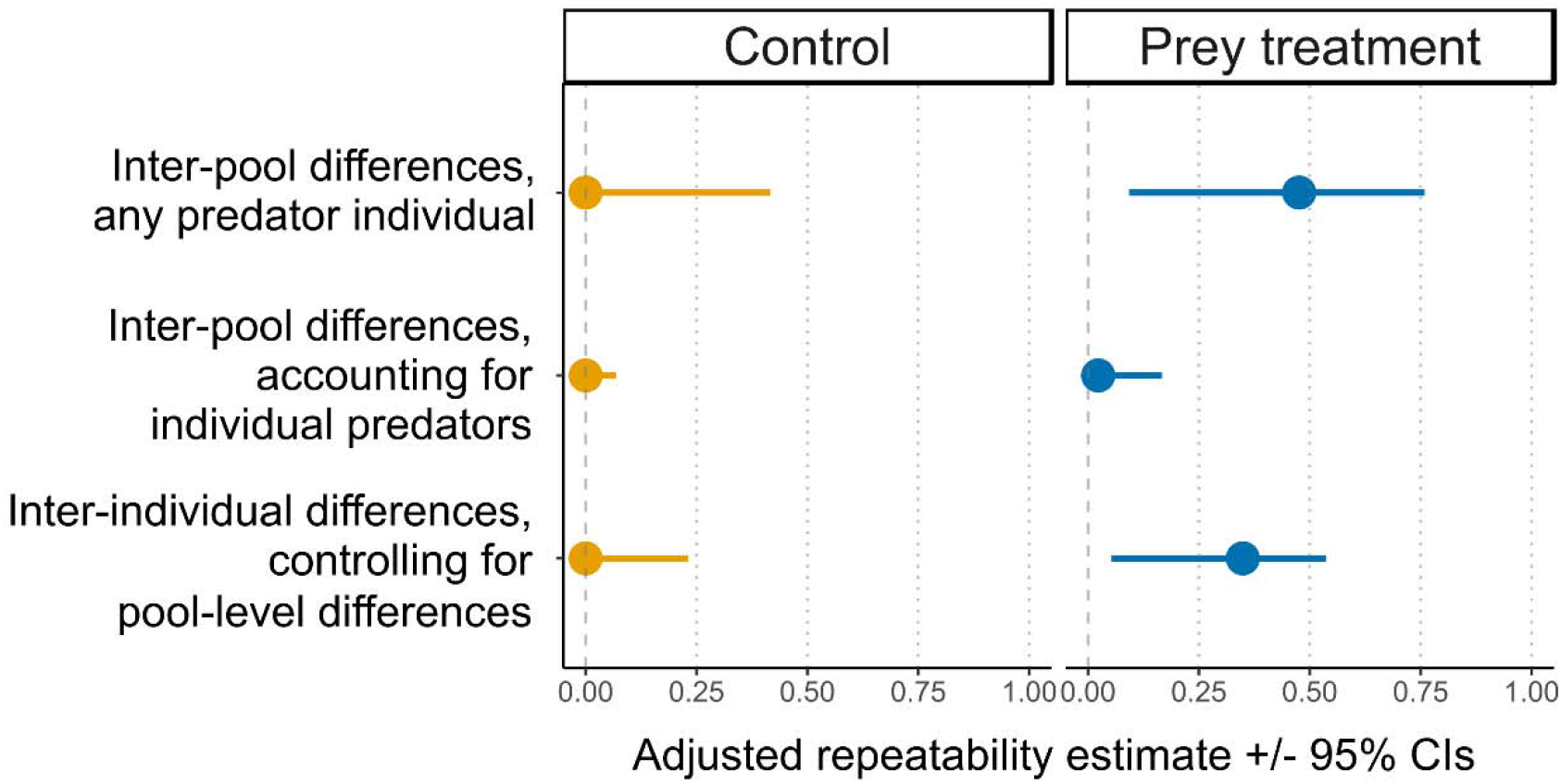
Pool- and individual-level repeatability estimates for the time spent near the stimulus, during control (orange) and prey treatment (blue) presentations. The figure shows adjusted repeatability estimates (filled circles), and the degree of uncertainty associated with each estimate (error bars representing 95% confidence intervals). Adjusted repeatabilities represent the proportion of the total variance in the time spent near the stimulus which can be attributed to variation between pools (top and middle rows) or individuals (bottom row), as opposed to variation within pools or individuals. Whereas estimates of inter-pool differences in the upper row are focused on the time spent near the stimulus by any predator individual (pool-level data, Appendix: Table A3), those in the middle and lower rows are based on analysis of the time individual predators spent near the stimulus (Appendix: Table A4).

### Variation between individual predators in their response to prey

To investigate inter-individual variation in predatory behaviour, adjusted repeatabilities were estimated for the time spent near the stimulus by individually-identified pike cichlids. This analysis was limited to pike cichlids which approached the stimulus in at least two separate prey treatment presentations, or in at least two separate control presentations. During prey treatment presentations, the time spent near the stimulus was significantly repeatable, even when controlling for standard body length and other experimental and environmental variables (*R*_ind_ = 0.349, 95% confidence interval: 0.053 - 0.537, *P* = 0.006, *N*_obs_ = 109, *N*_ind_ = 44; Fig. 3; Appendix: Table A4). In contrast, repeatable inter-individual differences were not observed in the control treatment (*R*_ind_ = 0, 95% confidence interval: 0 - 0.231, *P* = 0.5, *N*_obs_ = 59, *N*_ind_ = 24; Fig. 3; Appendix: Table A4). Whilst the time spent near the stimulus was positively correlated across prey treatment presentations, no correlations were evident across control presentations (Appendix: Fig. A2). Additionally, with the identity of individual predators factored into these analyses, there was no longer any substantial inter-pool variation in the time spent near the stimulus (*R*_pool_ = 0.023, 95% confidence interval: 0 - 0.167, *P* = 0.448, *N*_obs_ = 109, *N*_pool_ = 15; Appendix: Table A4). This suggests that the inter-pool differences resulted from variation in the behaviour of individual predators in separate pools, rather than other sources of inter-pool variation such as predator density or environmental factors.

### Possible effects of sampling bias and social interactions

Attempts to measure behaviour in natural populations are susceptible to bias because individuals vary in their tendency to engage with novel stimuli (Stuber et al. 2013), and because behavioural differences between individuals can be influenced by social interactions (McDonald et al. 2016). In our study, there was a positive association between the number of prey treatment presentations in which an individual approached the stimulus and the time it spent near the stimulus during the first prey presentation it was observed in (Poisson GLMM, *N*_obs_ = *N*_ind_ = 68: χ^2^_1_ = 10.983, *P* < 0.001; Appendix: Fig. A3 and Table A2, model 7). This implies that pike cichlids that approached the stimulus in multiple presentations were more predatory on average, relative to the overall predator population. The individual-level repeatabilities presented here (Fig. 3) may therefore under-estimate the true extent of inter-individual variation as they are likely to exclude the least predatory individuals. Randomisation simulations based on an approach developed by Ioannou et al. (2017) suggested that diversity in the response to prey was not dependent on the social environment within each pool (Fig. 4a). Additionally, there was no evidence for positive or negative interactions between predators within the same pool (Fig. 4b), suggesting that the observed inter-individual differences in predators’ response to prey were not magnified or suppressed as a result of social interactions.

**Figure 4:**
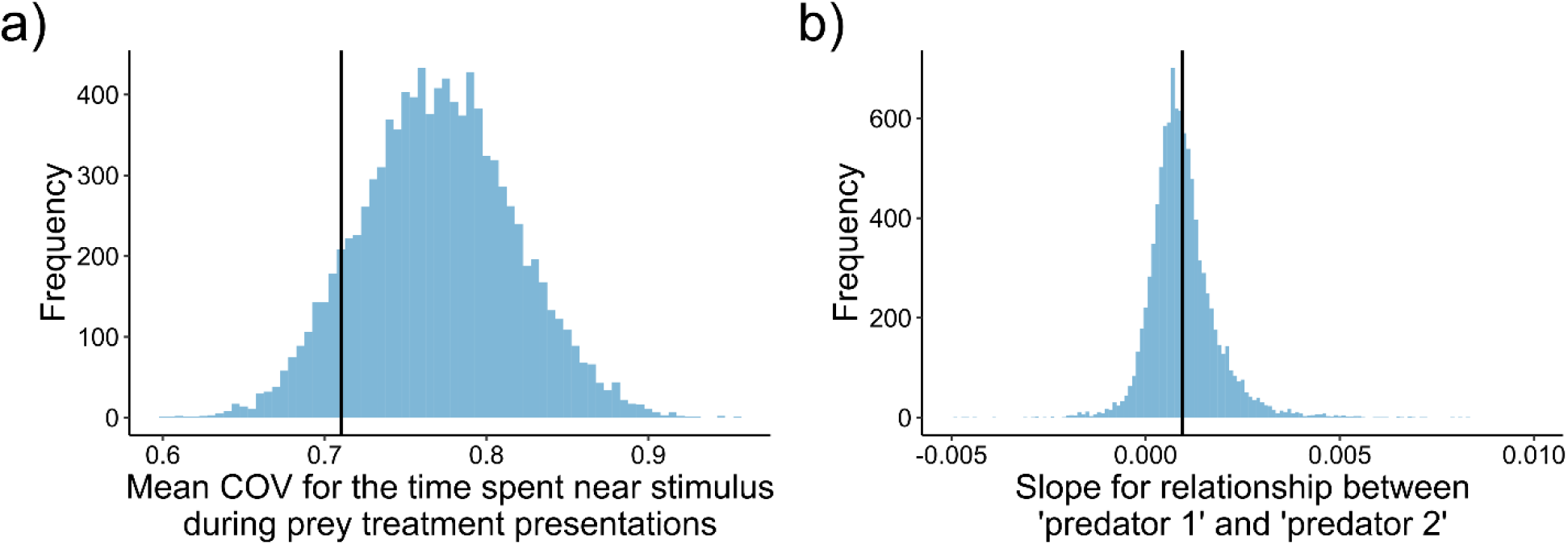
Results from randomisation simulations designed to examine the influence of social interactions on inter-individual variation in predators’ response to prey, based on 44 individual predators which approached the prey treatment stimulus in multiple presentations. a) The observed mean group-level diversity (COV) in predators’ responses to prey (indicated by the black vertical line) occurred within the 95% confidence intervals of the randomised distribution for the same statistic, produced when observations corresponding to individual predators were randomly shuffled between pools. b) The mean slope for the relationship between the time spent near the stimulus by pairs of fish randomly selected from the same pool was weakly positive (mean slope: 0.000976, shown by the black vertical line), but the 95% confidence intervals (−0.000431, 0.00299) of the randomised distribution of the slope overlapped with zero.

### Predicting individual predatory responses from behaviour when prey are absent

The comparison of treatments with and without prey shows that the behaviour of individual predators is only repeatable when prey are present. This suggests that the response to prey cannot be explained by personality variation in boldness or neophobia, as consistent differences arising from these traits should also lead to repeatability in the control presentations without prey. To test this explicitly, we explored whether the behaviour of pike cichlids during control treatment presentations could account for the repeatability in their response during prey treatment presentations. Adjusted repeatability estimates in the time spent near the stimulus during the prey treatment remained significantly repeatable when the mean time spent near the stimulus during control presentations was included as a covariate (*N*_obs_ = 109, *N*_ind_ = 44: *R*_ind_ = 0.33, 95% confidence interval: 0.013 - 0.506, *P* = 0.003). The repeatability estimate was still statistically significant when the time spent near the stimulus during the first control presentation an individual was observed in was instead included as a covariate (*N*_obs_ = 109, *N*_ind_ = 44: *R*_ind_ = 0.294, 95% confidence interval: 0.021 - 0.508, *P* = 0.006; in both analyses, individuals were assigned zero time spent near the stimulus in the control if they were not observed in the control treatments). These results are consistent with the absence of any correlations between the time spent near the stimulus during prey treatment presentations and the mean time spent near the stimulus across control presentations or time spent near the stimulus during the first control presentation in which an individual was observed (Appendix: Fig. A4 and Table A2: models 8-9). There was however a near-significant tendency for individuals observed in control presentations to spend more time near the prey treatment stimulus than those individuals which were never observed in the control (Appendix: Fig. A5 and Table A2, models 10-11), providing some indication of a role of boldness in determining the response of individual predators to their prey.

### Components of repeatability in predatory behaviour

One explanation for the increased repeatability of behaviour in the prey treatment compared to the control is that variation between individuals is greater in the presence of prey. To examine this possibility, we estimated inter-individual variances for the time spent near the stimulus during the control and prey treatments, which, unlike estimates of the repeatability, are not influenced by the consistency of individual behaviour (i.e. intra-individual variation) within each treatment. In contrast to negligible inter-individual variability among fish observed across control presentations (*N*_obs_ = 59, *N*_ind_ = 24, inter-individual variance: 0, 95% confidence interval: 0 - 0.353), inter-individual variability was apparent in the prey treatment (*N*_obs_ = 109, *N*_ind_ = 44, inter-individual variance: 0.597, 95% confidence interval: 0.064 - 0.977; Fig. 5). The residual variance was also higher in the control (1.645, 95% confidence interval: 0.844 - 2.118) compared to the prey treatment (1.074, 95% confidence interval: 0.713 - 1.49), suggesting that intra-individual variation was lower when prey were present. Thus, the significant repeatability between individuals in the prey treatment and the lack of repeatability in the control can be explained by individuals being more variable relative to one another and also behaving more consistently in the prey treatment.

**Figure 5:**
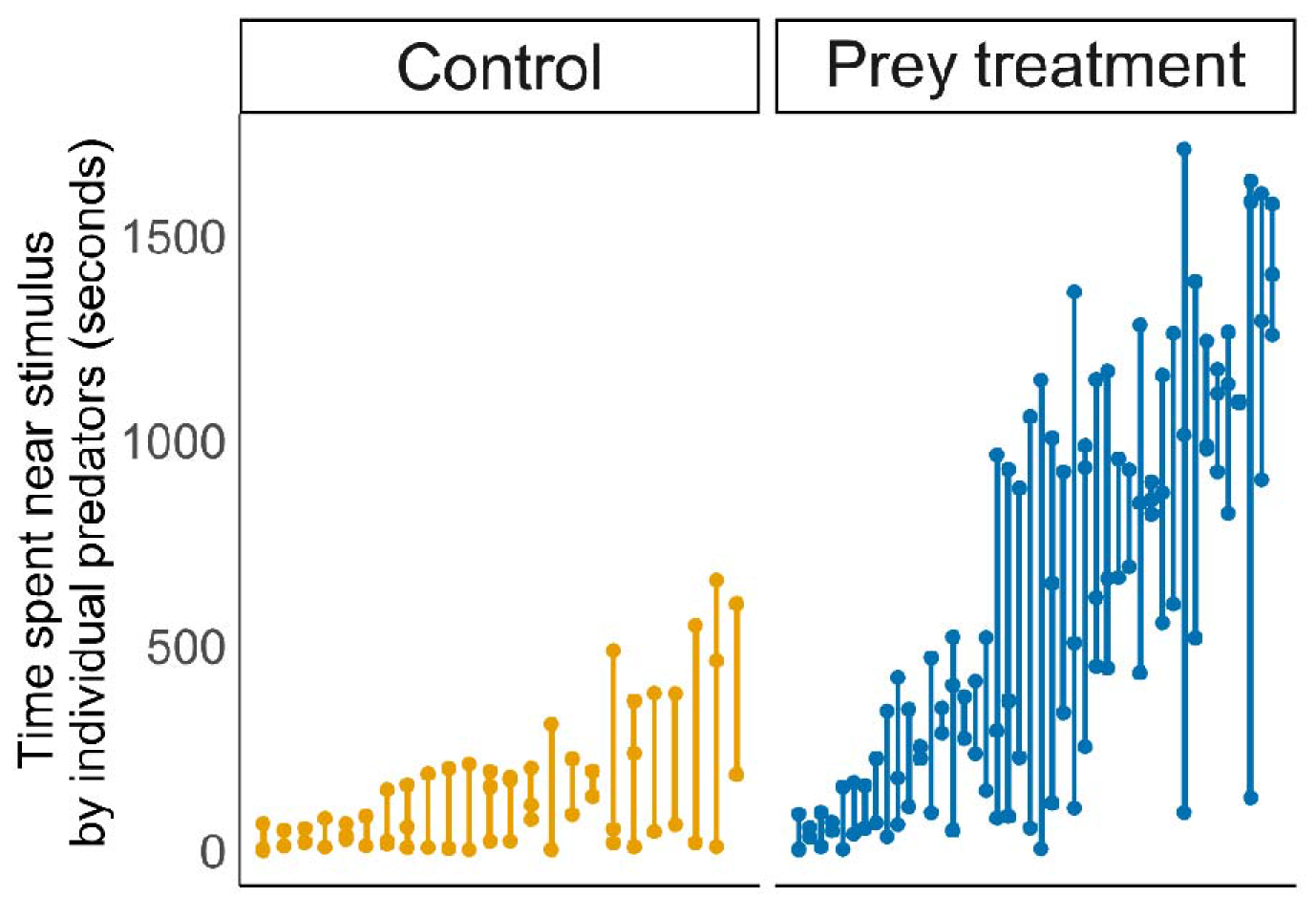
Variability in the behaviour of individual predators in the control (orange) and prey treatments (blue). Points indicate the time spent near the stimulus in each presentation, and the vertical lines span the range for each individual. Data is shown for individuals which approached the stimulus in two or more prey treatment presentations (*N*_ind_ = 44) or two or more control presentations (*N*_ind_ = 24), ordered by increasing mean time spent near the stimulus.

## Discussion

By studying the response of wild piscivorous fish to a stimulus prey shoal across a series of natural river pools, our study provides evidence for consistent inter-individual variation in the response of predators to their prey. After controlling for their body size and for experimental and environmental factors, we found that individual pike cichlids differed consistently in the time spent near the stimulus when prey were present, but not when prey were absent. Crucially, our analysis revealed that inter-individual variation in predatory behaviour could not be explained by the response of predators to the empty presentation apparatus (a novel object). This indicates that the response to prey reflects a personality trait which is specific to predation, is distinct from other commonly measured personality traits such as boldness or neophobia, and is independent of the previously documented correlation between boldness and encounter rates with prey (Pruitt et al. 2012). This individual-level trait accounted for variation between pools in the time spent near the stimulus by any pike cichlid, suggesting it is likely to explain a significant proportion of the risk faced by prey in their local environment, even when accounting for predator density. Our analysis also shows that inter-individual differences in the response to prey were unlikely to have arisen from pre-existing variation between predators becoming magnified due to the presence of prey, as boldness (as measured during control presentations) did not predict individual predators’ response to prey.

The existence of a personality trait specific to predation could affect prey in two main ways: by influencing the strength of non-lethal effects on prey traits, or by directly affecting survival during predator-prey encounters. Most obviously, the sustained or repeated presence of a nearby predator is likely to affect the level of risk perceived by prey. Prey often respond to the heightened risk of predation with a change in their behaviour, for example by increasing their vigilance, shifting microhabitat use or by forming social groups (Lima and Dill 1990). While anti-predator responses can be effective in enhancing survival over the short-term, these responses often reduce the amount of time allocated to foraging or reproductive behaviour, leading to long-term fitness costs. In many instances, the potency of non-lethal predator effects on prey fitness components such as individual growth rates, survival or reproduction can rival the impact of direct prey consumption (Preisser et al. 2005), with potentially substantial consequences for prey demographic rates (Eggers et al. 2006), prey population dynamics (Peckarsky et al. 2008), interactions between prey and their competitors (Werner 1991) and the abundance of resources at lower trophic levels (Suraci et al. 2016). The behavioural response of prey to predation risk will depend on their perception of the impending threat, which will be sensitive to how predators in the vicinity are behaving (Stankowich and Blumstein 2005). For prey, a predator personality trait should make the level of background predation risk perceived by prey more predictable, particularly in situations where the same predator and prey individuals encounter one another frequently, as is the case in relatively isolated pools within the guppy-pike cichlid system.

A predation-specific personality trait could have a strong influence on the eventual outcome of encounters with prey and help to maintain behavioural variation within the prey population (McGhee et al. 2013). While our study was not designed to explore the possible direct effects of predator personality on prey survival, our results suggest that inter-individual variation in the predatory behaviour measured here may have potentially lethal consequences for prey. Although we found a positive association between the time spent near the stimulus and the overall probability of attack by pike cichlids, this association might be expected if the number of predatory strikes simply grows linearly with time. However, the positive relationship between the time spent by predators near the stimulus and the initial rate of attack (during the first 30 seconds spent near the stimulus) also implies that predators which spend more time around prey are more motivated to attack when prey are first encountered, and thus pose a greater threat. Having established that individual predators consistently differ in their response to a standardised prey stimulus independently of boldness, future research could investigate the impact of these differences on dynamic interactions between predators and prey, in a setting where prey are unconstrained and both predators and prey are free to respond to cues from one another. This would address whether prey adjust their anti-predator behaviour in response to predation-specific personality variation. If carried out in a setting where prey were exposed to individual predators for an extended period, this could also help clarify the relative impact on prey survival of a predation-specific personality trait compared to variation between individual predators in commonly studied personality traits which affect encounter rates, such as boldness or activity.

We also found evidence that consistent inter-individual variation in the behaviour of the predator individuals present within each pool accounted for differences between pools in the time spent near the stimulus by any pike cichlid. Pool-level differences were not correlated with the densities of predators in each pool, suggesting that the personality traits of resident predators could be an important factor contributing to local differences in predation risk. However, in addition to predator density, pools might also differ in the social environment predators are exposed to. If the presence of an individual near the stimulus alerts others to the presence of prey through social information (Pitcher et al. 1982), inter-individual differences might be suppressed by social interactions. Alternatively, if socially dominant predators aggressively exclude subordinates from accessing prey, feedbacks resulting from differences in social dominance could also magnify inter-individual variation in the response to prey (Bergmüller and Taborsky 2010). It was therefore important to account for the possibility that variation between pools in the nature and strength of social interactions could generate the observed inter-individual differences in predator behaviour (McDonald et al. 2016). In our study, randomisation simulations showed that the degree of inter-individual variation in the time spent near prey was not dependent on the observed distribution of individual pike cichlids between pools. This suggests that there were no social interactions between individual predators that positively or negatively affected inter-individual variation. In other words, being present within a particular pool, and exposed to a particular set of other individuals with specific behavioural characteristics, did not have a strong influence on an individual predator’s response to prey. Although it is difficult to fully disentangle intrinsic inter-individual differences from environmental influences on behaviour without translocating animals and quantifying their behaviour in different contexts (Niemelä and Dingemanse 2017), these findings support the conclusion that stable differences in social interactions were unlikely to have played a major role in shaping the observed inter-individual variation in the response to prey.

By demonstrating that the risk prey experience cannot always be adequately predicted from frequently studied axes of personality variation, our study highlights the importance of considering inter-individual variation in traits with direct ecological relevance. These results also have specific implications for the guppy-pike cichlid system, in which geographically isolated guppy populations occupying low- and high-predation environments demonstrate dramatic differences in numerous aspects of their life history (Reznick et al. 1990) and behaviour (Ioannou et al. 2017). Since high-predation zones are characterised by the presence of pike cichlids, the existence of differences in predator behaviour between pools adds to the accumulating evidence that predation pressure is more heterogeneous within these areas than implied by the well-studied contrast between high- and low-predation environments (Deacon et al. 2018).

## Acknowledgments

We thank Robert Heathcote and Darren Croft for advice in the field, and thank Emmanuelle Briolat, Hannah MacGregor, Iestyn Penry-Williams and Teresa Szopa for helpful comments on the manuscript. This work was supported by a NERC GW4+ Doctoral Training Partnership studentship from the Natural Environment Research Council [NE/L002434/1] awarded to ASC, and a Natural Environment Research Council Fellowship [NE/K009370/1] and a Leverhulme Trust grant [RPG-2017-041 V] awarded to CCI.

