## Supplementary figures and tables for "Predatory behaviour as a personality trait in a wild fish population"

**Appendix for ‘Predatory behaviour as a personality trait in a wild fish population’**

**Figure A1:** Relationship between log-transformed time spent near the stimulus and presentation number, in control and prey treatment presentations.

**Figure A2:** Correlations between the time spent near the stimulus during different presentations.

**Figure A3:** Relationship between the number of prey treatment presentations in which individual pike cichlids approached the stimulus and the time spent near the stimulus during the first presentation that an individual was observed in.

**Figure A4.** The relationship between the time spent near the stimulus during control presentations without prey and the time near the prey treatment stimulus.

**Figure A5.** The relationship between whether or not individual predators were observed in any of the control presentations without prey (a), or in the first control presentation (b), and the time spent near the stimulus during prey treatment presentations when prey were present.

**Table A1.** Locations, habitat characteristics and numbers of individual pike cichlids observed in the river pools included in the study.

**Table A2.** Full statistical results for GLMMs.

**Table A3.** Adjusted repeatability estimates (*R*_pool_) indicating the extent of consistent inter-pool differences in the time spent near the stimulus, in both prey treatment and control presentations.

**Table A4.** Adjusted repeatability estimates indicating the extent of consistent inter-individual (*R*_ind_) and inter-pool (*R*_pool_) differences in the time spent near the stimulus, in both control and prey treatment presentations.

**Video A1.** Example video showing a pike cichlid approaching and attacking the prey treatment stimulus.

**Literature cited in Appendix.**

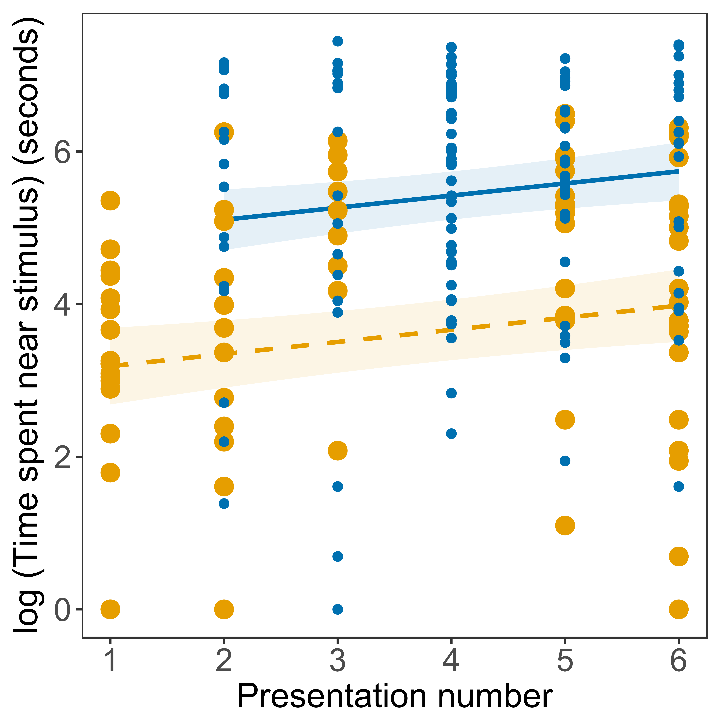

**Figure A1:** **Relationship between log-transformed time spent near the stimulus and presentation number, in control (large orange points, dashed orange line) and prey (small blue points, solid blue line) treatment presentations.** Shading represents 95% confidence intervals surrounding the predicted response derived from a GLMM featuring the time spent near the stimulus as the response (model 1 in table A2), with all other fixed effects not plotted above held constant at their mean values to obtain the predicted response. Pike cichlids spent more time near the stimulus with increasing presentation number (i.e., 1 to 6) (Poisson GLMM, *N*_obs_ (no. of observations) = 211, *N*_ind_ (no. of individuals): χ^2^_1_ = 7.30, *P* = 0.007), indicative of habituation to the apparatus and presentation procedure. The model was fitted to data at the level of a given presentation (211 observations) for all individual pike cichlids observed approaching the apparatus (69 individuals). The model included the following variables as fixed effects: stimulus treatment (control vs. prey), presentation number, time of day (to account for the effects of diurnal cycle on pike cichlid behaviour) (Endler 1987), canopy openness (levels of incident light may effect visibility and therefore predator-prey interactions) and the estimated number of pike cichlids in each pool (obtained from video analysis). Continuous fixed effects were also standardised by subtracting the mean and dividing by the standard deviation for each variable. Pool and individual identity were included as nested random intercepts, to account for non-independence arising from repeated measures of the same individuals and clustering of multiple individuals within each pool. An observation-level random intercept was also included in the model to counter over-dispersion (Harrison 2014).

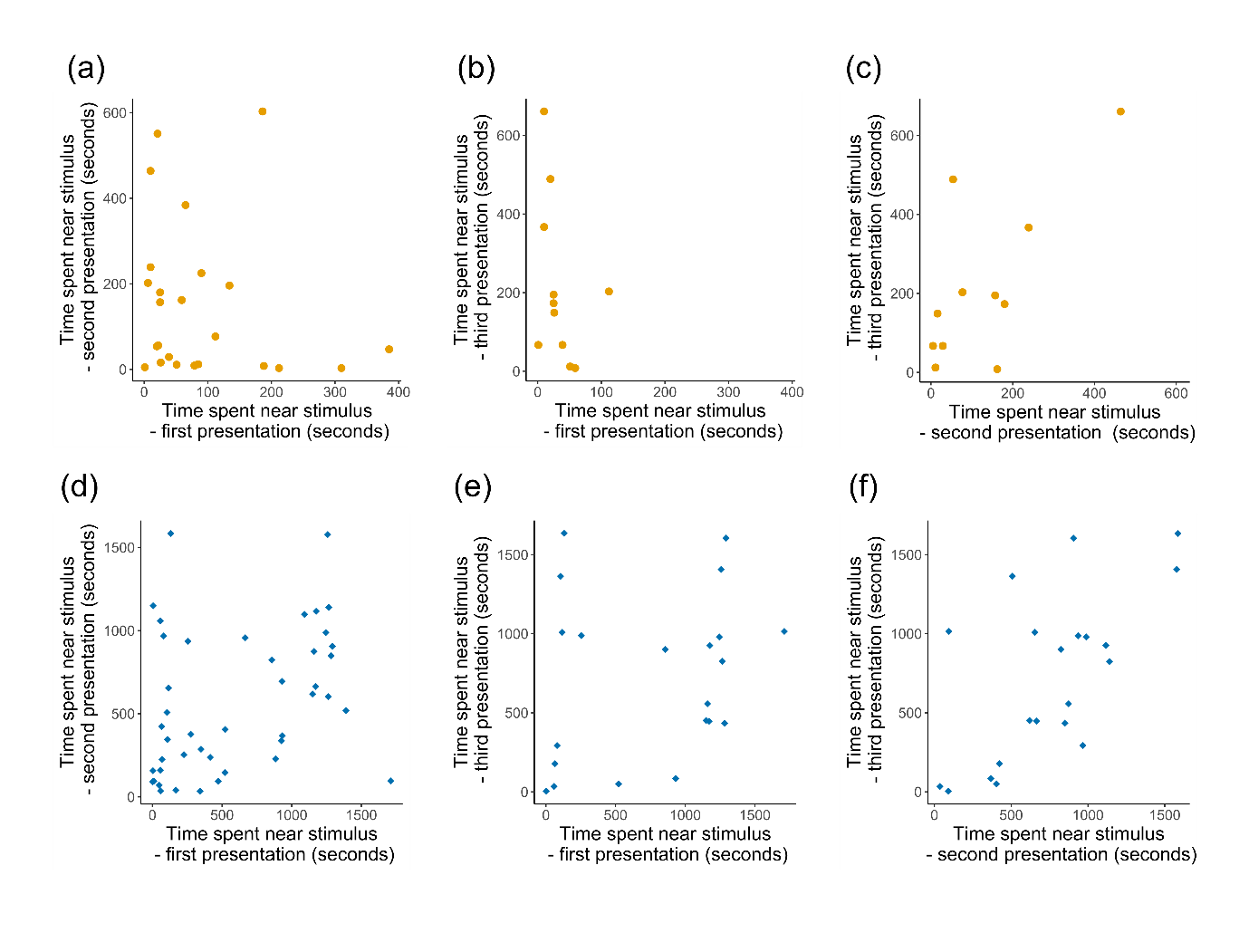

**Figure A2:** **Correlations between the time spent near the stimulus during different presentations.** Plots show comparisons between different control presentations ((a)-(c), orange dots) and prey treatment presentations ((d)-(f), blue diamonds). Shown are the first versus second presentations in which a pike cichlid was observed (a), (d), first and third presentations (b), (e), and the second and third presentations (c), (f).

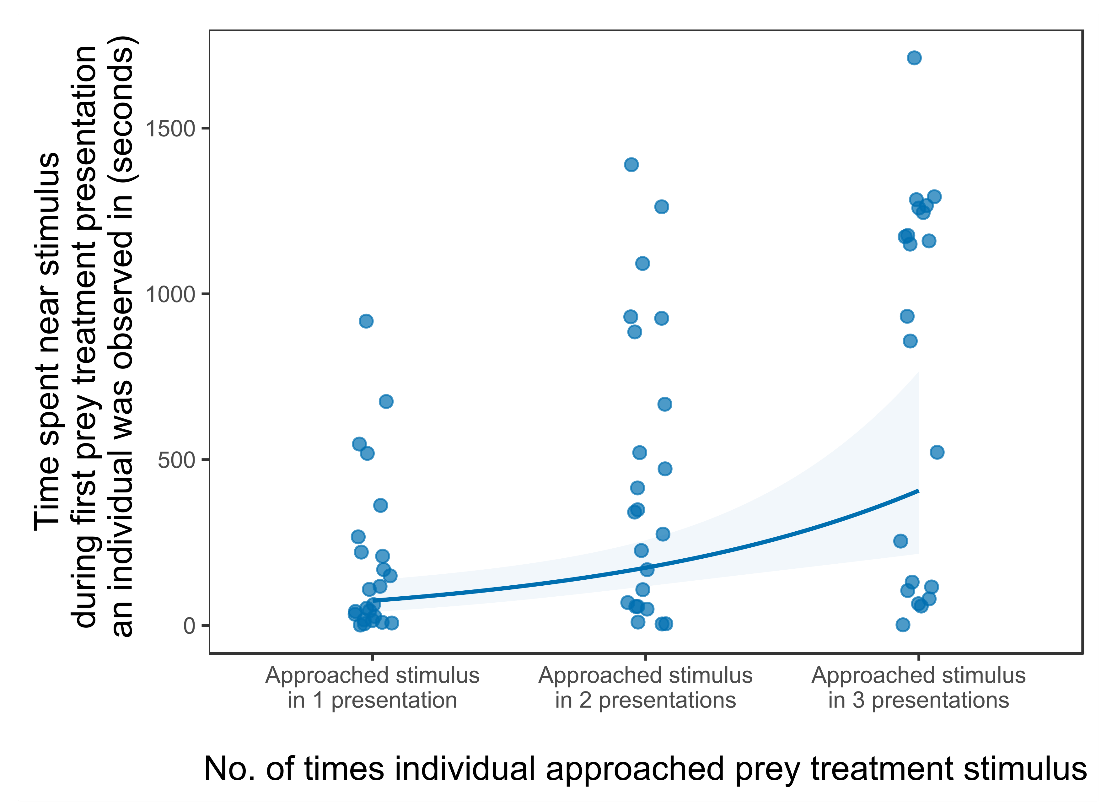

**Figure A3. Relationship between the number of prey treatment presentations in which individual pike cichlids approached the stimulus and the time spent near the stimulus during the first presentation that an individual was observed in.** Pike cichlids which approached the prey treatment stimulus during multiple presentations were more predatory on average, compared to those which approached in only one presentation. The curve represents the predicted response derived from a Poisson GLMM (model 7 in table A2) and the shading represents uncertainty (95% confidence intervals) surrounding this response. The model was fitted to data on the response of all individual pike cichlids observed interacting the stimulus during prey treatment presentations (68 individuals). Pool and individual identity were included in the model as nested random intercepts, and an observation-level random intercept term was also included to counter over-dispersion (Harrison 2014). In the above figure, data points are also offset laterally to increase visibility.

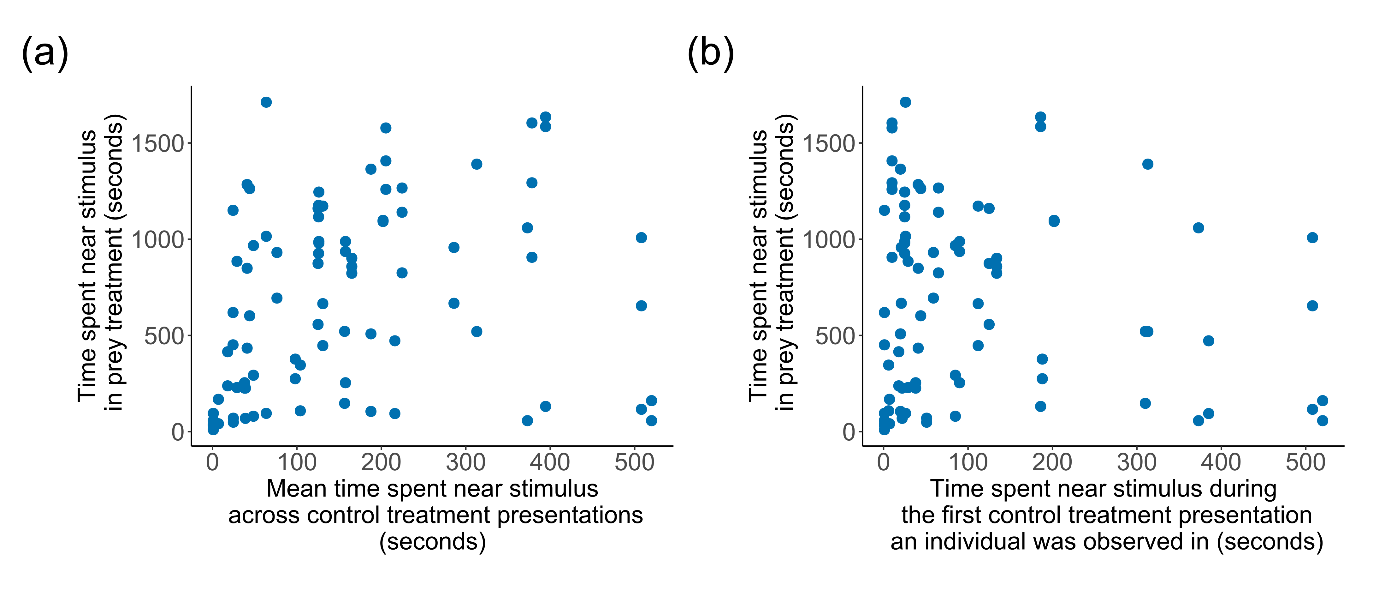

**Figure A4.** **The relationship between the time spent near the stimulus during control presentations without prey and the time near the prey treatment stimulus.** The time spent near the stimulus by individual predators in separate prey treatment presentations is plotted against the mean time spent near the stimulus across all three control presentations per pool (a), and the time spent near the stimulus during the first control presentation in which an individual was observed (b). Data is shown for individual predators which approached the stimulus in two or more prey treatment presentations and which were also observed approaching the stimulus in at least one control treatment presentation (*N*_ind_ = 35). There was no correlation between the time spent near the stimulus in the prey treatment and the mean amount of time spent near the stimulus over the three control presentations (Poisson GLMM, *N*_obs_ = 87, *N*_ind_ =35: χ^2^_1_ = 0.426, *P* = 0.514, model 8 in table A2), or with the time spent near the stimulus during the first control presentation in which an individual was observed (Poisson GLMM, *N*_obs_ = 87, *N*_ind_ = 35: χ^2^_1_ = 0.0695, *P* = 0.792, model 9 in table A2).

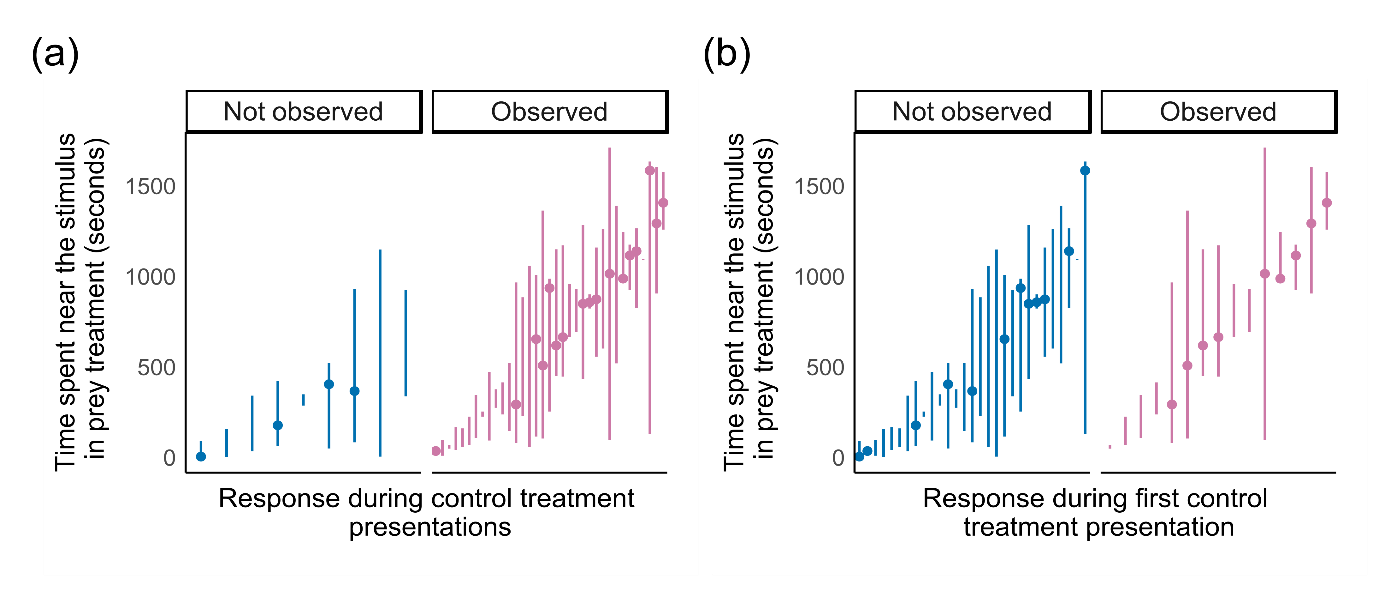
**Figure A5.** **The relationship between whether or not individual predators were observed in any of the control presentations without prey (a), or in the first control presentation (b), and the time spent near the stimulus during prey treatment presentations when prey were present.** In cases where individuals approached the stimulus in two prey treatment presentations, the vertical lines represent the maximum and minimum amount of time an individual spent near the stimulus in the two separate presentations. In cases where individuals approached the stimulus in three separate prey treatment presentations, the dots indicate the median value for the time spent near the stimulus. Data is shown for, and analyses were based on, individuals which approached the stimulus in two or more prey treatment presentations (*N*_ind_ = 44). There was a near-significant tendency for individual pike cichlids observed approaching the stimulus during at least one control presentation to spend more time near the stimulus during prey treatment presentations (a: Poisson GLMM, *N*_obs_ = 109, *N*_ind_ = 44: χ^2^_1_ = 3.17, *P* = 0.0749, model 10 in table A2). However, individuals which were observed in the first control presentation did not differ in the time spent near the stimulus during the prey treatment from those which were not observed in the initial presentation (b: Poisson GLMM, *N*_obs_ = 109, *N*_ind_ = 44: χ^2^_1_ = 1.14, *P* = 0.285, model 11 in table A2).

**Table A1.** **Locations, habitat characteristics and numbers of individual pike cichlids observed in the river pools included in the study.** C1/P overlap refers to individuals recorded over multiple prey treatment presentations and in at least one control presentation, or in multiple prey treatment presentations and multiple control presentations (C2/P overlap). Data from 16 pools were analysed, as in one of the pools (pool 13) no pike cichlids were observed approaching the stimulus, and in another pool (pool 9) there were a large number of pike cichlids which prevented individuals from being reliably identified.

| Pool | Location (latitude, longitude) | Canopy openness (%) | No. of individuals observed approaching the stimulus in either treatment | No. of individuals observed approaching the stimulus over multiple presentations | | | |
| --- | --- | --- | --- | --- | --- | --- | --- |
|  |  |  |  | Control, C | Prey treatment, P | C1/P overlap | C2/P overlap |
| 1 | 10⁰42.143’N, 61⁰19.239’W | 11.09 | 2 | 0 | 2 | 0 | 0 |
| 2 | 10⁰42.173’N, 61⁰19.277’W | 8.58 | 2 | 2 | 2 | 2 | 2 |
| 3 | 10⁰42.197’N, 61⁰19.284’W | 8.49 | 3 | 1 | 1 | 1 | 0 |
| 4 | 10⁰42.258’N, 61⁰19.277’W | 64.91 | 9 | 2 | 5 | 5 | 2 |
| 5 | 10⁰42.286’N, 61⁰19.256’W | 0.00 | 3 | 3 | 2 | 2 | 2 |
| 6 | 10⁰42.293’N, 61⁰19.246’W | 54.08 | 8 | 1 | 3 | 2 | 1 |
| 7 | 10⁰41.988’N, 61⁰19.239’W | 26.52 | 2 | 1 | 2 | 2 | 1 |
| 8 | 10⁰41.039’N, 61⁰19.277’W | 70.29 | 8 | 0 | 6 | 4 | 0 |
| 10 | 10⁰41.971’N, 61⁰19.252’W | 63.26 | 4 | 2 | 2 | 2 | 1 |
| 11 | 10⁰41.976’N, 61⁰19.269’W | 52.78 | 6 | 3 | 4 | 2 | 2 |
| 12 | 10⁰42.001’N, 61⁰19.262’W | 22.36 | 7 | 0 | 5 | 3 | 1 |
| 14 | 10⁰42.258’N, 61⁰19.277’W | 41.73 | 2 | 0 | 0 | 0 | 0 |
| 15 | 10⁰41.336’N, 61⁰19.487’W | 10.20 | 2 | 1 | 2 | 2 | 1 |
| 16 | 10⁰42.327’N, 61⁰19.243’W | 12.10 | 2 | 1 | 1 | 1 | 1 |
| 17 | 10⁰42.334’N, 61⁰19.240’W | 3.38 | 3 | 2 | 2 | 2 | 2 |
| 18 | 10⁰42.377’N, 61⁰19.210’W | 2.60 | 6 | 5 | 5 | 5 | 5 |
| **Totals** | | | **69** | **24** | **44** | **35** | **21** |

**Table A2.** **Full statistical results for GLMMs.** Results for Poisson GLMMs used to assess the influence of experimental and environmental variables on the time spent near the stimulus (models 1-2), binomial GLMMs used to assess the influence of the time spent near the stimulus by an individual pike cichlid on the probability of attacking the stimulus during the same presentation (models 3-4), Poisson GLMMs used to assess the relationship between the time spent near the stimulus (over a whole presentation) on the number of attacks made by pike cichlids during the first 30 seconds they spent near the stimulus (models 5-6) and a Poisson GLMM used to assess the relationship between the number of separate prey treatment presentations in which an individual pike cichlid was observed and the time spent near the stimulus during the first prey treatment presentation in which an individual was observed (model 7). Poisson GLMMs were also used to explore the relationship between the response of predators to prey and the behaviour of the same individuals in control presentations (models 8-11). Models 1-2, 3-4 and 5-6 differ in including either presentation number (i.e. 1 to 6, accounting for any habituation or learning effects; models 1, 3 and 5) or the proportion of previous prey treatment presentations (accounting for the sequence in which control and prey treatment presentations were presented; models 2, 4 and 6) as a fixed effect. Separate (but otherwise identical, in terms of the other fixed effects and random terms that were included) models were constructed in order to avoid problems associated with collinearity (calculation of variance inflation factors (VIFs) revealed collinearity between these two explanatory variables (VIF>3)) (Graham 2003, Zuur et al. 2010). The results remained similar whether presentation number or the proportion of previous prey treatment presentations were included in the model. All models included pool-level random intercepts, models 1-6 and 8-11 included individual-level random intercepts (nested within pools) and models 1-2 and 7-9 included an observation-level random intercept to counter over-dispersion (Harrison 2014). Sample sizes for the subset of data used to fit each model are given in the first column (*N*_obs_ = no. of observations, *N*_ind_ = no. of individuals).

| **Model** | **Response variable** | **R^2^_GLMM_** | **Explanatory variable(s)** | **Estimate** | **S.E.** | **t value** | ***P-*value** |
| --- | --- | --- | --- | --- | --- | --- | --- |
| Model 1 (Poisson GLMM) | Time spent near stimulus | 0.302 | Stimulus treatment | 1.759 | 0.192 | 9.173 | <0.001 |
| *N*_obs_ = 211  *N* _ind_ = 69 |  |  | Presentation number | 0.252 | 0.092 | 2.730 | 0.007 |
|  |  |  | Time of day | -0.248 | 0.148 | -1.681 | 0.098 |
|  |  |  | Canopy openness | -0.633 | 0.207 | -3.065 | 0.009 |
|  |  |  | Estimate number of pike cichlids per pool | 0.395 | 0.206 | 1.914 | 0.064 |
| Model 2 (Poisson GLMM)  *N*_obs_ = 211  *N* _ind_ = 69 | Time spent near stimulus | 0.306 | Stimulus treatment | 1.636 | 0.200 | 8.169 | <0.001 |
|  |  |  | Proportion of previous prey treatment presentations | 0.296 | 0.097 | 3.045 | 0.003 |
|  |  |  | Time of day | -0.304 | 0.146 | -2.085 | 0.041 |
|  |  |  | Canopy openness | -0.687 | 0.204 | -3.366 | 0.005 |
|  |  |  | Estimated number of pike cichlids per pool | 0.460 | 0.202 | 2.279 | 0.030 |
| Model 3 (binomial  GLMM)  *N*_obs_ = 133  *N* _ind_ = 68 | Probability of attack (binary variable indicating whether or not the stimulus was attacked during a presentation) | 0.886 | Time spent near stimulus | 6.847 | 2.043 | 3.352 | <0.001 |
|  |  |  | Standard body length | 1.290 | 0.435 | 2.966 | <0.001 |
|  |  |  | Presentation number | -0.433 | 0.337 | -1.286 | 0.188 |
|  |  |  | Time of day | 0.299 | 0.342 | 0.873 | 0.379 |
|  |  |  | Canopy openness | 0.620 | 0.589 | 1.052 | 0.291 |
|  |  |  | Estimated number of pike cichlids per pool | -0.394 | 0.582 | -0.677 | 0.496 |
| Model 4 (binomial GLMM)  *N*_obs_ = 133  *N* _ind_ = 68 | Probability of attack (binary variable indicating whether or not the stimulus was attacked during a presentation) | 0.899 | Time spent near stimulus | 7.327 | 2.303 | 3.182 | <0.001 |
|  |  |  | Standard body length | 1.296 | 0.433 | 2.991 | <0.001 |
|  |  |  | Proportion of previous prey treatment presentations | -0.443 | 0.354 | -1.250 | 0.198 |
|  |  |  | Time of day | 0.361 | 0.345 | 1.048 | 0.291 |
|  |  |  | Canopy openness | 0.621 | 0.582 | 1.068 | 0.285 |
|  |  |  | Estimated number of pike cichlids per pool | -0.440 | 0.581 | -0.757 | 0.445 |
| Model 5  (Poisson GLMM)  *N*_obs_ = 123  *N* _ind_ = 63 | Number of attacks in the first 30 seconds pike cichlids spent near the stimulus | 0.263 | Time spent near stimulus | 0.212 | 0.079 | 2.685 | 0.009 |
|  |  |  | Standard body length | 0.323 | 0.074 | 4.378 | <0.001 |
|  |  |  | Presentation number | -0.095 | 0.065 | -1.473 | 0.141 |
|  |  |  | Time of day | 0.152 | 0.138 | 1.101 | 0.274 |
|  |  |  | Canopy openness | -0.008 | 0.201 | -0.040 | 0.968 |
|  |  |  | Estimated number of pike cichlids per pool | 0.158 | 0.194 | 0.815 | 0.421 |
| Model 6  (Poisson GLMM)  *N*_obs_ = 123  *N* _ind_ = 63 | Number of attacks in the first 30 seconds pike cichlids spent near the stimulus | 0.260 | Time spent near stimulus | 0.200 | 0.079 | 2.533 | 0.014 |
|  |  |  | Standard body length | 0.331 | 0.074 | 4.466 | <0.001 |
|  |  |  | Proportion of previous prey treatment presentations | -0.120 | 0.069 | -1.746 | 0.085 |
|  |  |  | Time of day | 0.167 | 0.137 | 1.222 | 0.231 |
|  |  |  | Canopy openness | -0.008 | 0.198 | -0.041 | 0.966 |
|  |  |  | Estimated number of pike cichlids per pool | 0.138 | 0.191 | 0.716 | 0.475 |
| Model 7 (Poisson GLMM)  *N*_obs_ = 68  *N* _ind_ = 68 | Time spent near stimulus during first prey treatment presentation pike cichlid was observed in | 0.154 | Total number of prey treatment presentations an individual pike cichlid was observed in | 0.849 | 0.245 | 3.470 | <0.001 |
| Model 8 (Poisson GLMM)  *N*_obs_ = 87  *N* _ind_ = 35 | Time spent near the stimulus, prey treatment  presentations | 0.192 | Mean time spent near stimulus across all control presentations | 0.099 | 0.154 | 0.646 | 0.514 |
|  |  |  | Presentation number | 0.092 | 0.099 | 0.927 | 0.357 |
|  |  |  | Time of day | -0.096 | 0.180 | -0.531 | 0.597 |
|  |  |  | Canopy cover | -0.597 | 0.284 | -2.102 | 0.056 |
|  |  |  | Estimated number of pike cichlids per pool | 0.561 | 0.277 | 2.023 | 0.068 |
| Model 9 (Poisson GLMM) *N*_obs_ = 87  *N* _ind_ = 35 | Time spent near the stimulus, prey treatment presentations | 0.192 | Time spent near stimulus, during first control treatment presentation in which an individual was observed | -0.03574 | 0.136 | -0.264 | 0.792 |
|  |  |  | Presentation number | 0.0934 | 0.010 | 0.943 | 0.349 |
|  |  |  | Time of day | -0.136 | 0.185 | -0.738 | 0.466 |
|  |  |  | Canopy cover | -0.630 | 0.290 | -2.172 | 0.053 |
|  |  |  | Estimated number of pike cichlids per pool | 0.571 | 0.285 | 2.001 | 0.072 |
| Model 10 (Poisson GLMM)  *N*_obs_ = 109  *N* _ind_ = 44 | Time spent near the stimulus, prey treatment presentations | 0.206 | Whether or not individual was observed in any control presentations (binary variable) | 0.628 | 0.347 | 1.809 | 0.071 |
|  |  |  | Presentation number | 0.026 | 0.004 | 6.231 | <0.001 |
|  |  |  | Time of day | 0.371 | 0.018 | 20.886 | <0.001 |
|  |  |  | Canopy cover | -0.142 | 0.265 | -0.536 | 0.597 |
|  |  |  | Estimated number of pike cichlids per pool | 0.202 | 0.250 | 0.806 | 0.433 |
| Model 11 (Poisson GLMM)  *N*_obs_ = 109  *N* _ind_ = 44 | Time spent near the stimulus, prey treatment presentations | 0.197 | Whether or not individual was observed in the first control presentation (binary variable) | 0.319 | 0.297 | 1.074 | 0.285 |
|  |  |  | Presentation number | 0.026 | 0.004 | 6.236 | <0.001 |
|  |  |  | Time of day | 0.371 | 0.018 | 20.853 | <0.001 |
|  |  |  | Canopy cover | -0.175 | 0.273 | -0.641 | 0.530 |
|  |  |  | Estimated number of pike cichlids per pool | 0.220 | 0.259 | 0.849 | 0.412 |

**Table A3.** **Adjusted repeatability estimates (*R*_pool_) indicating the extent of consistent inter-pool differences in the time spent near the stimulus, in both prey treatment and control presentations.** The statistical significance of each estimate was assessed using a combination of P-values (obtained through likelihood ratio tests) and overlap of the 95% confidence intervals with zero (computed via parametric bootstrapping, denoted in square brackets). *N*_obs_ and *N*_pool_ refer to the number of observations and pools used in these analyses, respectively.

| Variables controlled for | Treatment | *N*_obs_ | *N*_pool_ | *R*_pool_ | *P* |
| --- | --- | --- | --- | --- | --- |
| time of day, presentation number | Control | 35 | 15 | 0.007 [0, 0.462] | 1 |
|  | Prey treatment | 45 | 16 | 0.605 [0.258, 0.828] | < 0.001 |
| time of day, presentation number, no. of pike cichlids in each pool, canopy openness | Control | 35 | 15 | 0 [0, 0.417] | 1 |
|  | Prey treatment | 45 | 16 | 0.476 [0.093, 0.759] | 0.009 |

**Table A4.** **Adjusted repeatability estimates indicating the extent of consistent inter-individual (*R*_ind_) and inter-pool (*R*_pool_) differences in the time spent near the stimulus, in both control and prey treatment presentations.** The statistical significance of each estimate was assessed using a combination of *P*-values (obtained through likelihood ratio tests) and overlap of the 95% confidence intervals with zero (computed via parametric bootstrapping, denoted in square brackets). *N*_obs_*, N_i_*_nd_ and *N*_pool_ respectively indicate the number of observations, individuals and pools included in these analyses. Adjusted repeatability estimates were similar regardless of whether presentation number (rows 1-2, table below) or the proportion of previous prey treatment presentations (rows 3-4, table below) were included as fixed effects (separate GLMMs were constructed to avoid problems associated with collinearity between these two variables) (Zuur et al. 2010).

| Variables controlled for | Stimulus treatment | *N_o_*_bs_ | *N*_ind_ | *R*_ind_ | *P* | *N*_pool_ | *R*_pool_ | *P* |
| --- | --- | --- | --- | --- | --- | --- | --- | --- |
| time of day, presentation number, no. of pike cichlids in each pool, canopy openness, standard body length | Control | 59 | 24 | 0 [0, 0.231] | 0.500 | 12 | 0 [0, 0.069] | 0.500 |
|  | Prey treatment | 109 | 44 | 0.349 [0.053, 0.537] | 0.006 | 15 | 0.023 [0, 0.167] | 0.448 |
| time of day, proportion of previous prey treatment presentations, no. of pike cichlids in each pool, canopy openness, standard body length | Control | 59 | 24 | 0 [0, 0.219] | 0.500 | 12 | 0 [0, 0.010] | 0.500 |
|  | Prey treatment | 109 | 44 | 0.322 [0.050, 0.519] | 0.007 | 15 | 0.049 [0, 0.190] | 0.374 |

**Video A1.** **Example video showing a pike cichlid approaching and attacking the prey treatment stimulus.**
